# Broad-Spectrum HIV-1 Detection and Neutralization via Multivalent Designer DNA Nanostructures

**DOI:** 10.1101/2025.09.21.677611

**Authors:** Saurabh Umrao, Abhisek Dwivedy, Mengxi Zheng, Varada Anirudhan, Ugur Parlatan, Timothy Henrich, Brian T. Cunningham, Utkan Demirci, Lijun Rong, Xing Wang

## Abstract

Early and accurate detection of HIV-1 remains a critical unmet need, particularly during the acute phase of infection when viral loads are low and transmission risk is highest. Here, we report a modular diagnostic and antiviral platform based on designer DNA nanostructures engineered for high-affinity recognition of the HIV-1 envelope glycoprotein (GP120). A custom DNA aptamer, termed HINT, was developed to bind GP120 across major HIV-1 subtypes (Groups M and P; subtypes A and B) with nanomolar affinity. To amplify binding strength, HINT aptamers were spatially patterned onto a net-shaped DNA nanostructure (DNA-Net_HINT_) that geometrically matches the trimeric GP120 spikes on the viral surface. Using multivalent interactions, the nanostructure enabled up to 10^4^-fold improvement in binding affinity (sub-picomolar *K*_*D*_), confirmed by surface plasmon resonance. Integration of DNA-Net_HINT_ into a paper-based lateral flow assay produced a low-cost, saliva-compatible self-testing device capable of detecting intact HIV-1 virions at concentrations as low as 328 viral copies per test, outperforming commercial fourth-generation rapid diagnostic tests. In addition to its diagnostic capabilities, the DNA-Net_HINT_ construct exhibited potent antiviral activity, reducing pseudovirus infection with an EC_50_ of ∼1.8 nM, nearly 1,000-fold more effective than free aptamers. This work demonstrates a dual-function DNA nanotechnology platform that enables both ultrasensitive HIV-1 detection and entry inhibition. The approach is broadly applicable to other enveloped viruses and represents a promising step toward next-generation molecular theranostics for infectious disease management.

## Introduction

Early and accurate detection of human immunodeficiency virus type 1 (HIV-1) remains critical for curbing viral transmission and improving clinical outcomes^1^. Despite significant advances in diagnostic technologies, current molecular tests rely predominantly on the amplification and detection of viral genetic material or viral antigens, such as HIV RNA and the p24 protein^2^. HIV RNA typically becomes detectable in plasma approximately 10 days post-infection, whereas the p24 antigen emerges 4–10 days subsequently, detectable via fourth-generation rapid diagnostic tests (RDTs), including the Determine™ HIV-1/2 Ag/Ab Combo (Abbott) kits^3^. However, both RNA and antigen detection methods are impeded by intrinsic limitations. Specifically, the evolving antibody response during early infection results in the formation of immune complexes with p24, significantly reducing antigen detection sensitivity and extending the “window period” for diagnosis from several weeks to months^4,5^. Critically, this window period coincides with a markedly elevated transmission risk, accounting for up to 50% of new HIV infections^6^. Moreover, existing assays exhibit compromised sensitivity due to HIV-1 genetic diversity and rapid antigenic drift, further limiting their effectiveness^7,8^. Thus, there is an unmet need for assays capable of highly sensitive, selective, and rapid detection of intact HIV-1 virions across diverse viral strains during early stages of infection.

The HIV-1 virion is composed of spike trimeric envelope glycoproteins (GP120), which serve as the sole virally encoded antigens on the viral surface. These proteins facilitate viral entry into host cells by binding to CD4 receptors and are organized into sparse trimeric complexes (∼14 per virion), presenting conserved epitopes accessible for direct virion detection^9,10^. This structural and functional specificity makes GP120 an ideal candidate for targeted detection assays. Synthetic single-stranded DNA (ssDNA) aptamers, designed to mimic antibody functionality, present a promising solution for sensitive and specific GP120 targeting, due to their ease of chemical synthesis, high specificity, and robust affinity for their targets^11,12^. Leveraging proprietary intellectual property (IP) tools, we developed an <u>HI</u>V-1 D<u>N</u>A aptamer candidate, termed “HINT.” While the specific aptamer selection strategy and sequence details are withheld due to patent filings, we conducted a comparative assessment against the previously reported HD4 aptamer, known for its diagnostic utility in detecting HIV-1 GP120^13-15^. The HINT aptamer demonstrated markedly improved binding affinity and broader target recognition. Notably, it binds with high affinity across HIV-1 group M and group P, as well as subtypes A and B, thereby addressing a critical limitation of existing aptamer-based assays in the context of viral diversity. These findings were supported by computational modeling studies and experimental surface plasmon resonance (SPR) data. The HINT aptamer was strategically designed to address two critical challenges in early HIV diagnostics: (1) the direct detection of intact virions in unprocessed saliva samples, thereby enhancing diagnostic sensitivity and reducing false-negative rates during acute infection^16^; and (2) robust recognition of GP120 antigens across diverse HIV-1 strains to maintain diagnostic accuracy despite viral antigenic drift.

To address the limited avidity inherent in monovalent aptamer-GP120 interactions, exacerbated by the low density and wide spatial distribution of GP120 trimers on the viral envelope^17^, we employed DNA nanotechnology to engineer a multivalent binding platform^18^. This approach is particularly relevant given HIV-1’s well-characterized immune evasion mechanism, whereby the sparse arrangement of envelope GP120 spikes (∼15 nm inter-spike spacing) prevents effective bivalent engagement by conventional immunoglobulin G (IgG) antibodies. The spatial separation exceeds the reach of IgG Fab arms, thereby limiting simultaneous binding to adjacent GP120 proteins and diminishing the overall efficacy of both immune recognition and therapeutic neutralization^19,20^.

To circumvent this geometric limitation, we devised a strategy in which HIV-targeting DNA aptamers were spatially organized into tri-aptamer clusters on a net-shaped DNA nanostructure, (henceforth referred as “DNA-Net”), mimicking the spatial topology of trimeric GP120 proteins on the virion surface^21^. The DNA-Net was designed using the multi-layer Designer DNA Nanostructure (DDN) framework, which enables programmable assembly of synthetic DNA strands into precisely defined nanoscale architectures with tunable shape, valency, spatial geometry, and surface functionality^21,22^. The structural bendability of the DNA-Net might enable a dynamic clustering of GP120 proteins to provide a maximal GP120 packing density and promoting avidity-driven multivalent interactions^9^. Analogous to DNA scaffold-guided clustering of cell surface mobile proteins, clustering of GP120 proteins guided by the DNA-Net scaffold can achieve global tri-aptamer–GP120 spatial pattern matching by correcting deviations of inter-GP120 distance through the multiple sites of attachment, resulting in a highly enhanced affinity of virus binding.

We subsequently translated the DNA-Net-based viral probes and assays into a rapid diagnostic tool based on lateral flow assay (LFA) principles. This resulted in the creation of a versatile, direct viral recognition platform capable of delivering rapid diagnostic results within 10 mins. The platform is highly accessible, with a per-test cost of approximately $1.20, substantially lower than the ∼$38 cost of commercial fourth-generation RDTs. Notably, our assay operates without the need for complex chemistry, signal amplification, or sample preprocessing, and it achieves detection sensitivity comparable to FDA-approved molecular methods. In clinical sample testing, the device successfully detected viral loads as low as 328 copies/mL, whereas commercial RDTs failed to detect viral loads below 10^5^ copies/mL under similar conditions.

In addition to diagnostic utility, flow cytometry-based assays demonstrated that DNA-Net viral probes significantly inhibit HIV-1 entry into host cells, with over 50% reduction in infection at nanomolar concentrations. These findings suggest dual functionality for the DNA-Net platform— as both a highly sensitive diagnostic tool and a potential antiviral agent. Taken together, our study highlights the translational potential of DNA nanostructure-based strategies for future theranostic applications in HIV-1 infection, offering a powerful and cost-effective platform for early detection and viral inhibition^23,24^.

## Results and Discussion

### Nanoscale molecular dynamics simulations for aptamer-GP120_MB_ protein interactions

The selection process for the HINT aptamer was developed using our proprietary tools. Due to patent constraints, we are unable to disclose the specific details of the selection process and the sequence information of the aptamer. To evaluate its binding characteristics, we performed a comparative analysis against the previously reported HD4 aptamer^25^, which was originally selected in silico for the GP120_MB_ (HIV-1 group M, subtype-B). We used the High Ambiguity Driven protein-protein DOCKing (HADDOCK) tool to predict the aptamer-bound protein complexes and Nanoscale Molecular Dynamics (NAMD) simulations for further refining the docked biomolecular complex quality^26,27^. Specifically, we examined the binding modes of individual monomeric aptamers, HINT/HD4 (**Supplementary Figure S1**), in comparison with a tri-aptamer configuration interacting with their corresponding GP120_MB_ protein targets (**Supplementary Figure S2**). Our objective was to determine whether the trimeric configuration could potentially foster more robust molecular interactions, leading to an enhanced binding affinity.

Monomeric aptamer configurations demonstrated a range of binding strengths, with interaction energies varying from relatively modest (–13.06 kJ/mol for HD4) to significantly stronger interactions (–63.60 kJ/mol for HINT) against single GP120_MB_ protomers **(Supplementary Figures S3-S4; Supplementary Movies S1-S2)**. Notably, the trimeric aptamer configurations exhibited substantially enhanced binding affinities: HD4 trimer complexes showed moderately increased binding energy (–39.94 kJ/mol), whereas HINT trimers presented very strong interactions, reaching –225.10 kJ/mol **(Supplementary Figure S5; Supplementary Movies S3-S4)**. The optimal trimeric aptamer complexes exhibited ∼6 nm spacing between aptamer-spike binding domains (**Supplementary Figures S6**), which closely matches with the native spacing of GP120 spikes on the HIV-1 envelope, as reported earlier^17,28^.

To gain further insight into the structural stability of the aptamer–GP120_MB_ protein complexes, we evaluated their conformational convergence during 192 ns of molecular dynamics simulations using pairwise Root Mean Square Deviation (RMSD) analysis. A plateau in RMSD values, or a region of minimal deviation across successive frames, is indicative of conformational convergence, signifying that the aptamer–protein complex has reached a relatively stable energy minimum within the simulation time scale. Such convergence often correlates with stronger intermolecular interactions, reduced structural fluctuations, and thermodynamic favorability of the binding interface. The HINT aptamer–GP120_MB_ trimeric complex achieves conformational convergence significantly earlier (around frame 1,000, ∼96 ns at 0.096 ns/frame) compared to the HD4–GP120_MB_ trimeric complex **(Supplementary Figure S5)**. This early convergence signifies rapid structural stabilization and extensive conformational rearrangements, resulting in the formation of a highly stable and energetically favorable HINT-protein complex **(Supplementary Figure S5B-S5C)**. Conversely, the HD4-protein complex exhibited delayed convergence and comparatively higher conformational variability, indicative of weaker intermolecular interactions and less optimal spatial complementarity with GP120 trimers **(Supplementary Figure S5E-S5F)**. This behavior is consistent with the deeper binding energy minimum observed in the trimeric configuration of HINT, further supporting its superior binding stability.

Further structural analysis of aptamer–protein interfaces revealed distinct differences in binding footprints between the two aptamers **(Supplementary Figures S2)**. The HD4 aptamer exhibited a preference for conserved (constant) regions of the GP120_MB_ protein, such as the C1 and C2 domains, in both its monomeric and trimeric configurations **(Supplementary Tables S1)**. While this targeting of invariant regions can ensure specificity, it may limit adaptability across antigenically diverse strains. In contrast, the HINT aptamer demonstrated a more versatile binding mode, engaging not only conserved regions but also multiple hypervariable loops, particularly V2, V3, and V4 across the GP120 subunits **(Supplementary Table S2)**. This expanded epitope recognition suggests enhanced conformational flexibility and a greater potential for cross-strain adaptability. Moreover, the HINT aptamer’s ability to interact with variable protein regions may allow it to better accommodate mutations and structural variations resulting from antigenic drift. Collectively, our molecular modeling and simulation data demonstrate that the HINT aptamer exhibits superior binding strength, structural stability, and epitope versatility compared to HD4.

### Surface plasmon resonance analysis of aptamer–GP120 protein interactions

To experimentally validate the computational predictions of enhanced HINT binding affinity, we employed surface plasmon resonance (SPR) to assess the binding interactions between aptamers and various subtypes and groups of HIV proteins (illustrated in **Figure 1a**). We focused first on HIV-1 group M proteins, which account for over 99% of global HIV infections^29^. Specifically, we analyzed aptamer binding to GP120 proteins from subtype A (GP120_MA_) and subtype B (GP120_MB_). Additionally, to evaluate cross-group reactivity, we tested the aptamers against GP120 from the genetically distinct HIV-1 group P (GP120_P_).

**Figure 1.**
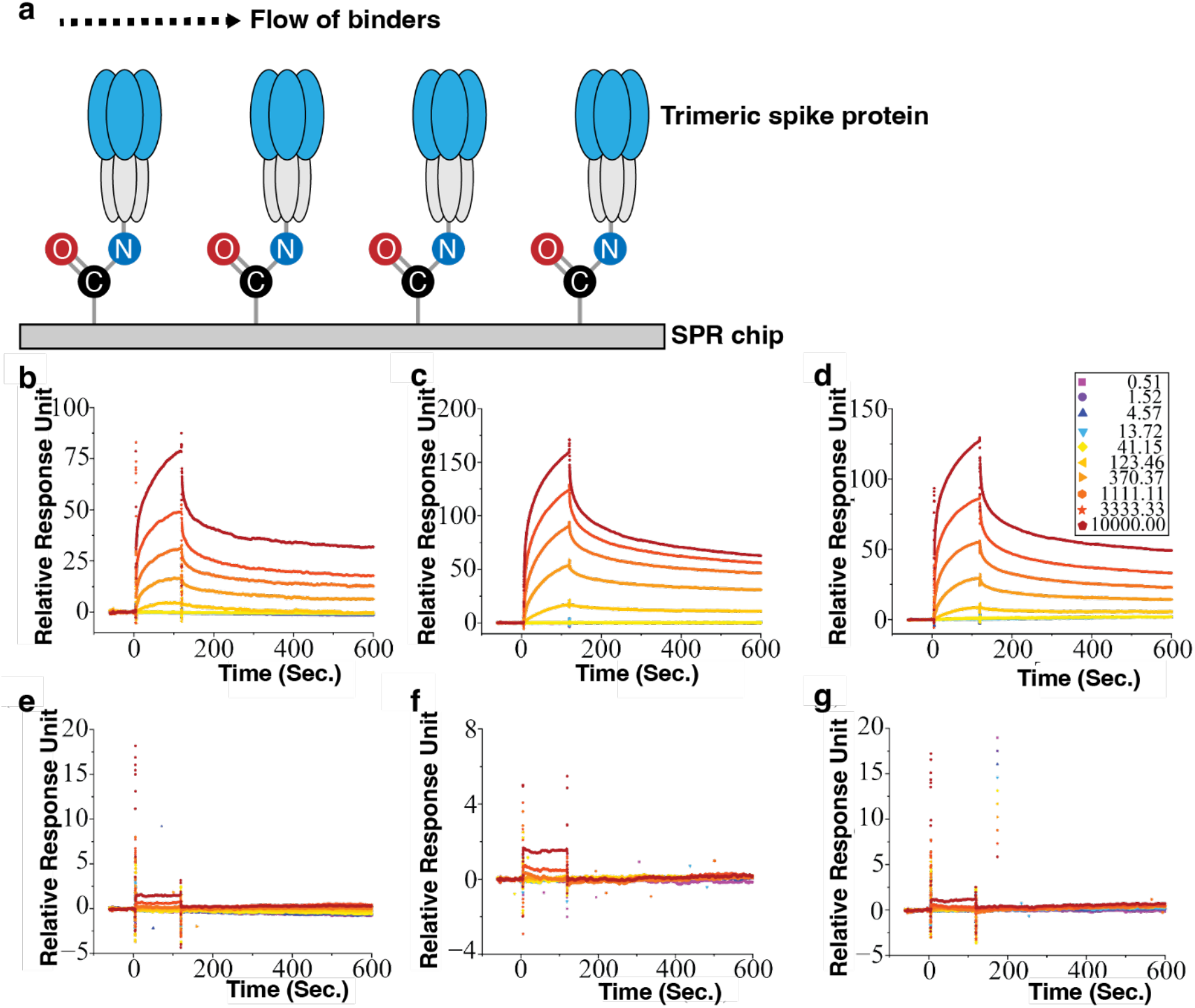
Surface plasmon resonance (SPR) assay. (**a**) Schematic of the SPR assay used to determine affinity of the aptamer binding. IgG protein at a concentration of 50 µg/mL served as the negative control in Flow Cell 1 (FC_1_). Panels **b** & **e**: Flow Cell 2 (FC_2_) was dedicated to immobilizing 50 µg/mL of HIV-1 Group M, subtype A isolate 92RW020 spike protein (GP120_MA_). Panels **c** & **f**: Flow Cell 3 (FC_3_) was employed to immobilize 50 µg/mL of HIV-1 Group M, subtype B (GP120_MB_). Panels **d** & **g**: Flow Cell 4 (FC_4_) was utilized to immobilize 50 µg/mL of HIV-1 Group P, strain RBF168 (GP120_P_). Subsequently, varying concentrations of aptamers (ranging from 0.51 nM –10000 nM) were introduced into each flow cell sequentially (FC_2_-FC_4_), for 600 s at a flow rate of 30 µL/min. Panels **b-d** show interaction of HINT aptamer with respective proteins present on FC_1_-FC_4_, while Panels **e-g** highlight the interaction of the HD4 aptamer with the corresponding proteins on FC_1_-FC_4_. Each sensorgram is repeated three independent times with similar results and corrected for non-specific interaction of aptamers with negative control. The resultant sensorgrams were employed to determine binding kinetics parameters, including the association rate constant (k_a_), dissociation rate constant (k_d_), and binding equilibrium dissociation constant (*K*_*D*_; *K*_*D*_ = k_d_/k_a_). These parameters were determined through global fitting of the complete association and dissociation phases using the 1:1 Langmuir binding model within BiaEvaluation software 4.0.1, developed by GE Healthcare Uppsala, Sweden.

SPR sensorgram analysis revealed that the HINT aptamer exhibits strong nanomolar-range binding affinities for all three GP120 proteins, with dissociation constants (*K*_*D*_) of 11.09 nM for GP120_MA_, 42.03 nM for GP120_MB_, and 80.67 nM for GP120_P_ **(Figures 1b–d)**. In contrast, HD4 showed minimal binding responses, particularly at lower concentrations, and failed to yield reliable *K*_*D*_ estimates across all tested variants **(Figures 1e–g)**. These experimental results corroborate the simulation findings and confirm that the HINT aptamer binds GP120 proteins from both dominant HIV-1 subtypes and across divergent viral groups with substantially higher affinity and breadth than HD4. This cross-clade reactivity suggests the HINT aptamer’s potential as a robust molecular recognition candidate for broad-spectrum HIV diagnostics and targeted antiviral strategies.

### DNA-Net design, synthesis, and characterization

Building upon our previous research, we implemented a “three-layer” DDN design principle to construct rhombus-shaped units of a DNA-Net structure, approximately measuring 45 nm × 45 nm^21,30^. Within this structure, we strategically arranged spike-targeting aptamers (comprising HINT/HD4) into an array of tri-aptamer clusters with a ∼6 nm intra-trimer spacing and ∼15 nm inter-trimer spacing to form DNA-Net_HINT_, and DNA-Net_HD4_ constructs **(Figure 2a)**. The intra-trimer cluster spacing, which provides first-layer pattern matching-based aptamer™spike interactions was adopted from the NAMD simulation data, where we found that the trimeric-aptamer configuration in a protein-bound complex maintains ∼ 6 nm intra-trimer spacing (**Supplementary Figure S6**). Furthermore, the 15 nm inter-tri aptamer spacing on the DNA-Net represents the smallest allowable spacing between two adjacent GP120 on the viral surface and thus, maximizing simultaneous inter-spike interaction with the tri-aptamers on the DNA-Net^17,28^. The resultant pattern-matching guided multivalent interactions provide an optimal framework for rapid virus binding with high avidity as employed in the build of LFA device in this study.

**Figure 2.**
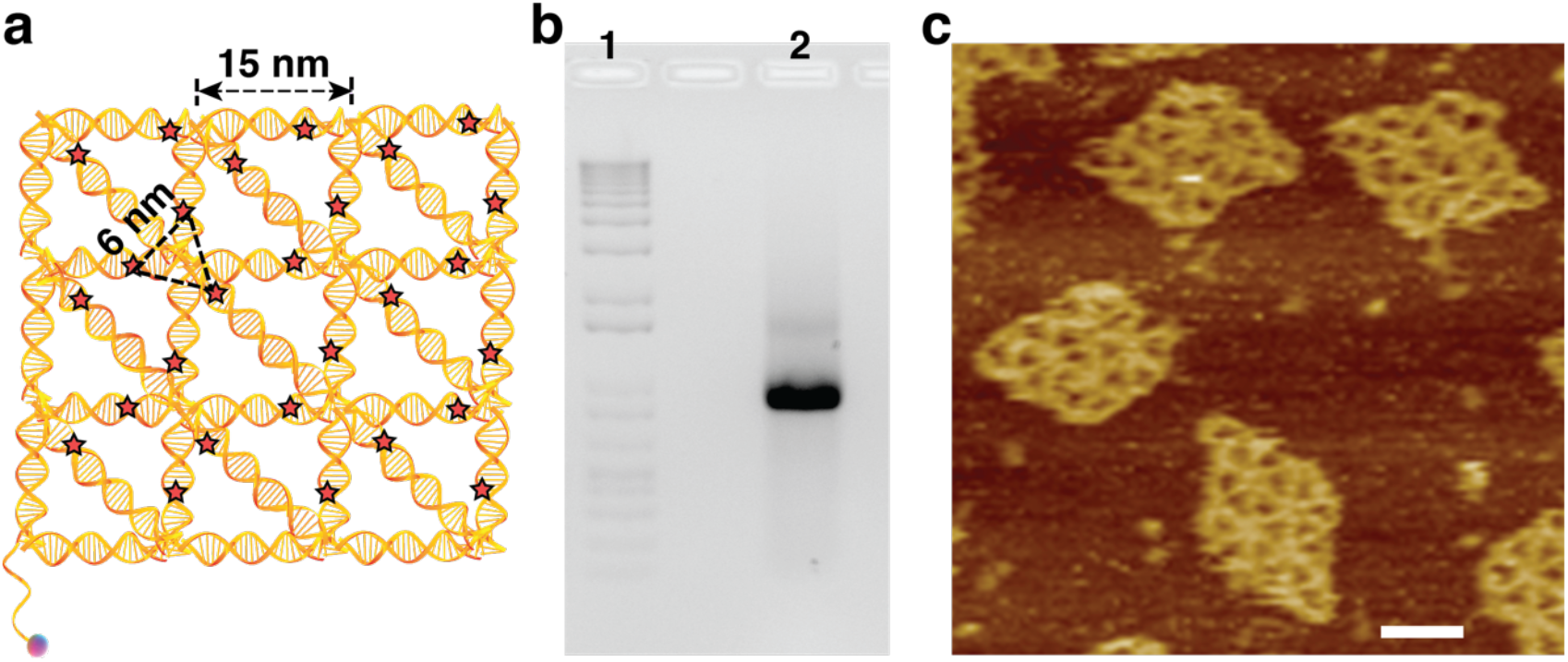
DNA-Net design, synthesis, and characterization. (**a**) Design and operation of the DNA-Net-based lateral flow assay (LFA) device. A rhombus-shaped DNA-Net, measuring 45 nm x 45 nm, was developed for this purpose. The ★ symbols indicate the positions of DNA aptamer clusters with a total of 27 tri-aptamer clusters distributed across the DNA-Net. Tri-aptamer clusters were strategically placed with approximately 6 nm intra-tri-aptamer spacing (as predicted by molecular dynamics simulations, **Supplementary Figure S6**) and 15 nm inter-tri-aptamer spacing,^18,28^ resembling the arrangement of spike trimers on a viral particle. (**b**) Formation of DNA-Net was analyzed using 1% agarose gel electrophoresis. (**c**) Atomic force microscopy (AFM) was employed in capturing high-resolution images of DNA-Net structure. Scale bar in the AFM image denotes a length of 25 nm.

The successful assembly of DNA-Net was first verified using 1% agarose gel electrophoresis (AGE, **Figure 2b**). Subsequently, the structural formation of the DNA-Net was confirmed using atomic force microscopy (AFM) imaging (**Figure 2c** and **Supplementary Figure S7**). A quantitative assessment of DNA-Net yield was performed using the “Gel Analysis” tool within ImageJ software, resulting in an approximate yield of 88% of DNA-Net monomer. The high purity of the desired DNA-Net structures obviated the necessity for additional purification steps. While the AGE analysis did reveal the presence of DNA-Net multimers in trace amounts, it is important to note that this minor amount would not exert a significant influence on the overall sensing capabilities of the DNA-Nets, as previously reported^21^.

### Surface plasmon resonance study with aptamer coupled DNA-Net and GP120 proteins

To evaluate the impact of multivalent presentation on aptamer binding performance, we conducted SPR assays comparing free aptamers with their DNA-Net-coupled counterparts. Specifically, we investigated the binding affinities of HINT and HD4 aptamers when presented in trimeric configurations on a DNA-Net scaffold (referred to as DNA-Net_HINT_ and DNA-Net_HD4_, respectively) against recombinant GP120 proteins from diverse HIV-1 subtypes and groups. Compared to the monomeric HINT aptamer **(Figure 1b–d)**, DNA-Net_HINT_ exhibited *K*_*D*_ in the picomolar range: 50.81 pM for GP120_MA_; 0.71 pM for GP120_MB_; and 7.48 pM for GP120_P_, representing a 200-fold (**Figure 3a**), a ∼60,000-fold (**Figure 3b)** and a ∼10,000-fold (**Figure 3c**), enhancement in affinity, respectively. These improvements are attributable to strong avidity effects and spatial pattern matching between the DNA-Net tri-aptamer clusters and the GP120 trimeric arrangement on the virion surface.

**Figure 3.**
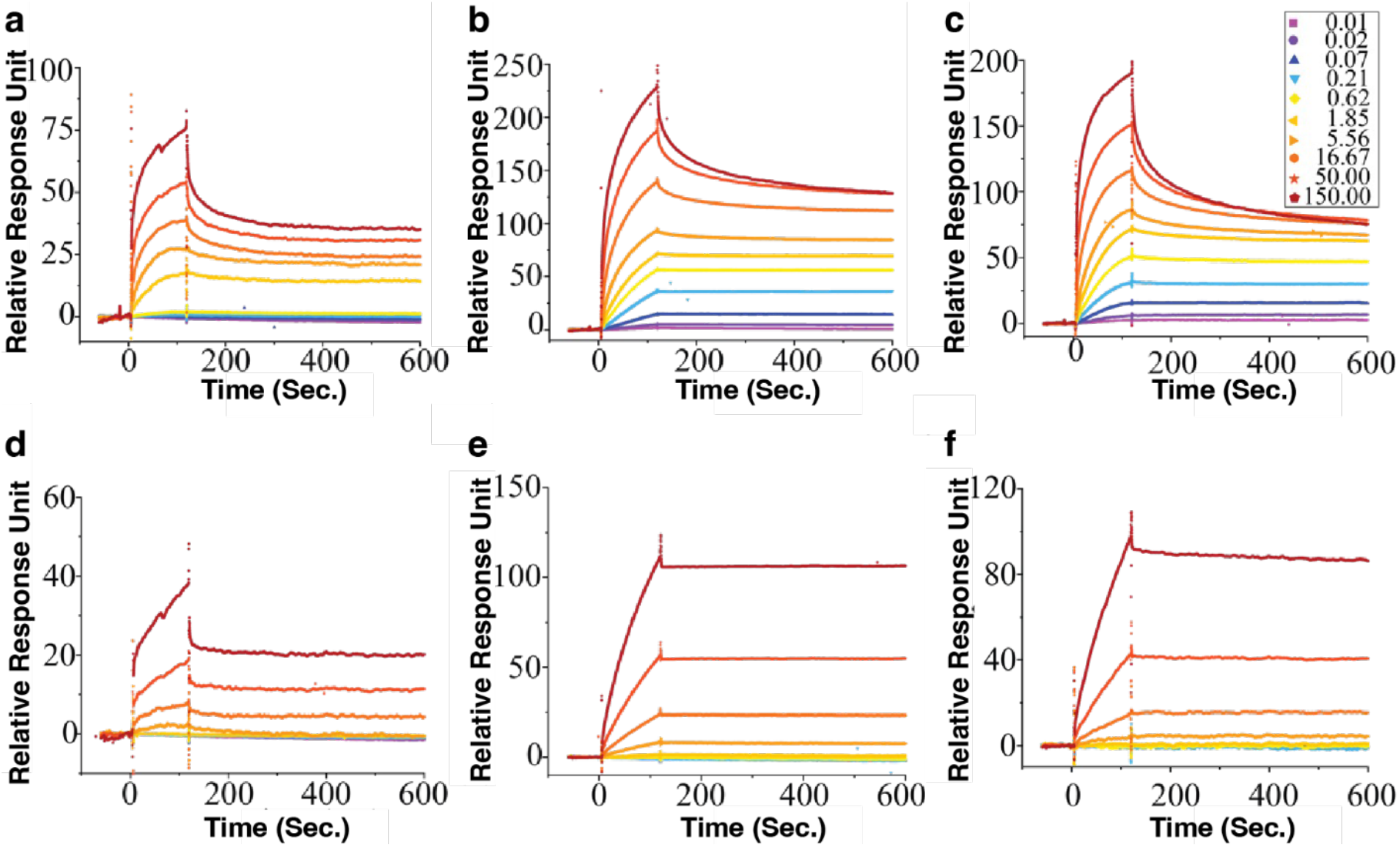
Surface plasmon resonance (SPR) assay. (**a**) Schematic of the SPR assay used to determine affinity of the DNA-Net-aptamer complex binding. IgG protein at a concentration of 50 µg/mL served as the negative control in Flow Cell 1 (FC_1_). Panels **a** & **d**: Flow Cell 2 (FC_2_) was dedicated to immobilizing 50 µg/mL of HIV-1 Group M, subtype A (GP120_MA_). Panels **b** & **e**: Flow Cell 3 (FC_3_) was employed to immobilize 50 µg/mL of HIV-1 Group M, subtype B (GP120_MB_). Panels **c** & **f**: Flow Cell 4 (FC4) was utilized to immobilize 50 µg/mL of HIV-1 Group P (GP120_P_). Subsequently, various concentrations of DNA-Net tethered with aptamers (0.01 nM –150 nM) were introduced into the respective flow cells for 600 s at a flow rate of 30 µL/min. Panels **a-c** show interaction of DNA-Net_HINT_ assays with respective proteins present on FC_1_-FC_4_, while Panels **d-f** highlight the interaction of the DNA-Net_HD4_ assay with the corresponding proteins on FC_1_-FC_4_. Each sensorgram is presented from three independent repetitions, yielding consistent results and corrected for the non-specific interaction of aptamers with the negative control.

Interestingly, even the HD4 aptamer, which showed weak or undetectable binding in its free monomeric form **(Figure 1e–g)**, exhibited measurable nanomolar affinity when incorporated into the DNA-Net structure. DNA-Net_HD4_ displayed *K*_*D*_ values of 1.03 nM (GP120_MA_), 0.10 nM (GP120_MB_), and 1.31 nM (GP120_P_) **(Figures 3d–f)**. These enhancements likely result from cooperative binding effects and increased local concentration of aptamers at the binding interface. The collective interaction of multiple HD4 aptamer units may partially compensate for their intrinsically lower monovalent affinity, enabling effective engagement with the GP120 trimers.

Collectively, these results validate the efficacy of our DNA-Net design in significantly enhancing the binding performance of aptamer-based molecular probes. The superior performance of DNA-Net_HINT_, in particular, showcases its selection as the optimal candidate for subsequent development of LFA-based HIV self-testing platform.

### Reporter assays formulation

The first step in the development of a LFA device is the formulation of a robust reporter (detector) assay. For this purpose, we utilized commercially available streptavidin-coated gold nanoshells (SA-AuNS), which are well known for generating strong and visually discernible signals in LFA-based platforms^31^. To construct the reporter assay, DNA-Net constructs (either DNA-Net_HINT_ or DNA-Net_HD4_) were conjugated to the SA-AuNS via biotin–streptavidin chemistry, resulting in the formation of AuNS_DNA-Net-HINT_ and AuNS_DNA-Net-HD4_ complexes. We performed a comprehensive physicochemical characterization to confirm successful conjugation and assess the stability of the nanoconjugates. Dynamic light scattering analysis revealed a marked increase in hydrodynamic diameter from 161.74 ± 2.50 nm (unmodified SA-AuNS) to 261.44 ± 1.91 nm upon conjugation with the DNA-Nets **(Supplementary Figure S8A)**. This size increase is consistent with theoretical predictions based on the dimensions of the DNA-Net constructs and indicates effective surface coverage of the nanoshells. It is important to note that the observed size distribution may vary depending on the geometric orientation of the DNA-Net scaffold on the nanoshell surface. In parallel, zeta potential measurements demonstrated a substantial shift in surface charge, with the initial value of −15.22 ± 0.44 mV for bare SA-AuNS decreasing to −35.4 ± 1.13 mV following DNA-Net conjugation **(Supplementary Figure S8B)**. This increase in negative surface charge reflects enhanced colloidal stability, attributed to the densely packed and uniformly distributed DNA-Net constructs on the nanoshell surface. These results validate the successful formulation of structurally stable and optically responsive reporter assays.

### Evaluation of DNA-Net -based LFA device using GP120_MB_ as target antigen

Guided by the favorable binding affinities observed in the SPR studies, we evaluated the sensitivity of our LFA device using the GP120_MB_ protein as a target. For this purpose, the DNA-Net_HINT_ construct was immobilized on the test line (denoted as DNA-Net_HINTatT_), while GP120_MB_ protein was immobilized on the control line, as capture probes, to enable sandwich complex formation and signal readout (**Figure 4a** for the convenience of illustration). A series of GP120_MB_ concentrations ranging from 1.95 nM to 250 nM was introduced in running buffer containing the AuNS_DNA-Net-HINT_ reporter conjugate. In the presence of target antigen, the DNA-Net_HINTatT_ and the AuNS_DNA-Net-HINT_ conjugate collectively forms a sandwich complex with the soluble GP120_MB_ on the test line (T). Concurrently, excess AuNS_DNA-Net-HINT_ forms a separate complex with the immobilized GP120_MB_ on the control line (C), ensuring proper device function. This interaction results in amplified gray-scale signal intensity on both the T and C lines, with the degree of signal correlating with increasing GP120_MB_ concentrations **(Figure 4b)**. The enhanced signal intensity enables both qualitative visual detection and semi-quantitative signal interpretation based on gray-scale intensity. Quantitative analysis revealed a broad linear detection range between 1.95 nM and 62.5 nM, with an apparent equilibrium dissociation constant (*K*_*D*_) of 62.97 ± 9.46 nM **(Figure 4c)**. The DNA-Net–based lateral flow assay demonstrates reliable and sensitive detection of free GP120_MB_ protein, supporting its potential applicability for detecting intact HIV-1 virions in diagnostic settings.

**Figure 4.**
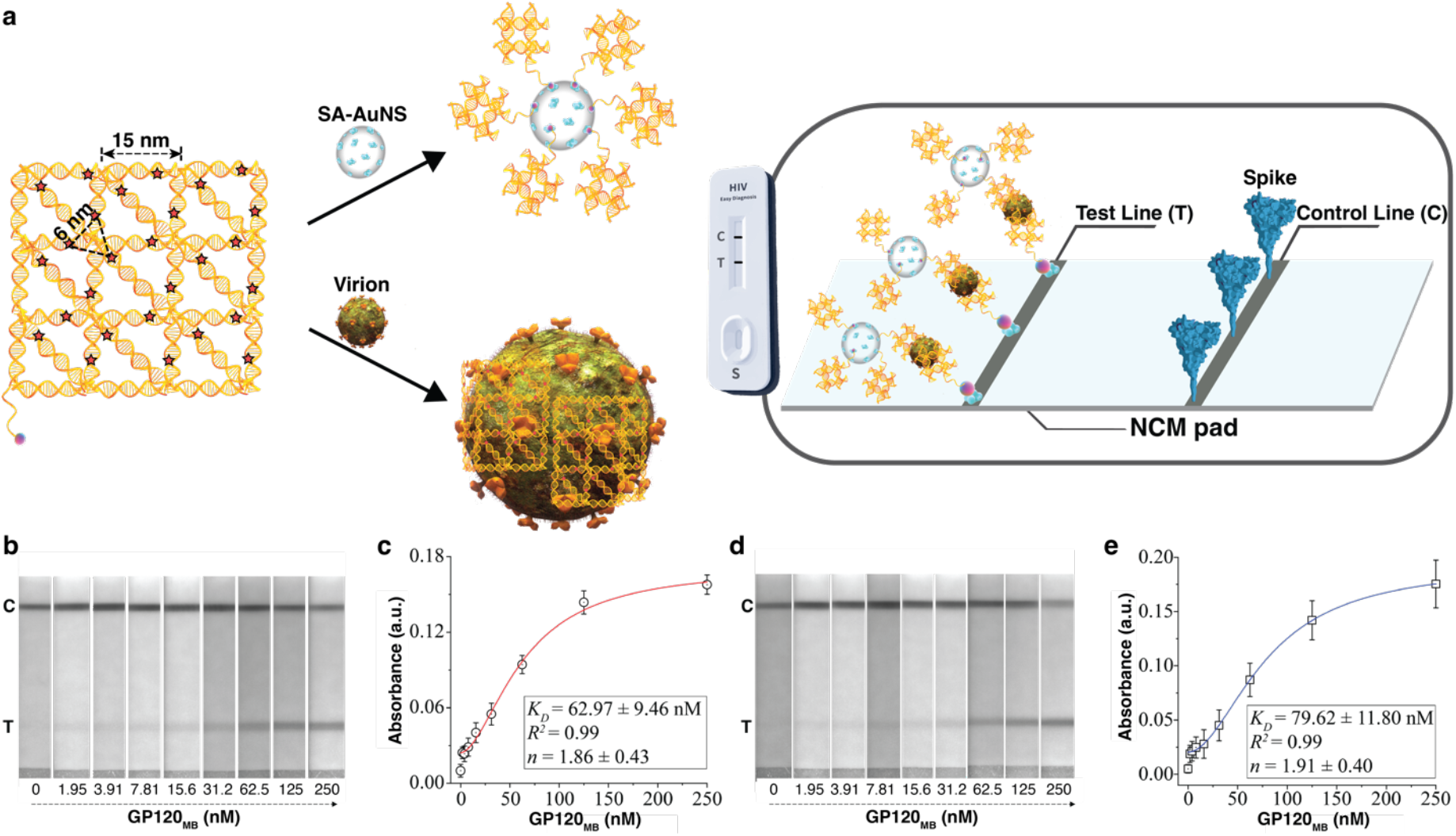
DNA-Net_HINT_ assay-based LFA device testing with GP120_MB_ protein as a target. (**a**) DNA-Net-based lateral flow assay (LFA) device is designed for the detection of spike proteins in free solution (**b, d** in this figure) and eventually on virion outer surface (**a-d** in **Figure 5**). A rhombus-shaped DNA-Net was developed for this purpose. The ★ symbols indicate the positions of DNA aptamer clusters with a total of 27 tri-aptamer clusters distributed across the DNA-Net. Tri-aptamer clusters were strategically placed with approximately 6 nm intra-tri-aptamer spacing and 15 nm inter-tri-aptamer spacing. Streptavidin-coated gold nanoshells (SA-AuNS) were labeled with DNA-Nets and employed as a reporter assay. In the LFA setup, 10 nM DNA-Net was applied to the test line using biotin-streptavidin chemistry, while 0.25 mg/mL of GP120 trimeric spike protein was deposited on the control line. Upon introducing spike protein or virus particles into the LFA device, they bind in a sandwich format with the reporter assay and DNA-Net located on the test line. Simultaneously, the reporter assay interacts with the GP120 on the control line through aptamer-protein interactions. The aggregation of AuNS leads to an enhancement in colorimetric signals observed in both the test and control lines. (**b**) The LFA device was subjected to a range of GP120_MB_ concentrations spanning from 1.95 nM to 250 nM, resulting in a gradual increase in the intensity of the test line (**T**). (**c**) Absorbance values from the test line were calculated and plotted against the respective GP120_MB_ concentrations. The binding isotherm of GP120_MB_ was analyzed using Hill fit. (**d**) The LFA device’s performance was assessed in a 50% saliva environment, with a similar range of GP120_MB_ concentrations. (**e**) Absorbance values from the test line were calculated and plotted. The binding isotherms of DNA-Net_HINT_ - GP120_MB_ were analyzed using Hill fit.

### Performance evaluation of DNA-Net-based LFA in biological fluids and specificity analysis

Given that rapid diagnostic tests for HIV screening are typically conducted using blood, oral fluid, and occasionally urine samples^32^, we evaluated the compatibility and robustness of our DNA-Net–based LFA device in clinically relevant biological matrices. To this end, we introduced a fixed concentration of 100 nM GP120_MB_ protein into solutions containing varying proportions (25% to 100%) of saliva, urine, and serum **(Supplementary Figure S9)**. The LFA device reliably detected the target antigen in media containing up to 75% saliva or urine. However, beyond 50% serum, false-positive signals were observed, while saliva and urine samples beyond this concentration threshold yielded false-negative results. These signal aberrations are likely attributable to high concentrations of endogenous proteins that interfere with the sandwich complex formation between the DNA-Net_HINTatT_ capture probe, the GP120_MB_ target, and the AuNS_DNA-Net-HINT_ reporter conjugate. Additionally, increased fluid viscosity at higher biological fluid concentrations may impede capillary flow within the LFA matrix, thereby compromising assay kinetics and reducing signal resolution. Based on these observations, a 50% saliva composition was identified as the optimal working condition, offering a balance between physiological relevance and device performance.

Under this optimized condition, we evaluated the binding characteristics of the DNA-Net_HINT_ assay using GP120_MB_ concentrations ranging from 1.95 nM to 250 nM in a 50% saliva matrix **(Figure 4d)**. The resulting binding isotherm revealed a broad linear detection range of 1.95 nM to 62.5 nM, with a *K*_*D*_ of 79.62 ± 11.80 nM **(Figure 4e)**, which closely aligned with the *K*_*D*_ obtained under aqueous conditions **(Figure 4c)**. The minimal shift in affinity suggests that diluted salivary proteins exert only modest influence on target recognition, thereby affirming the assay’s robustness in biologically complex environments.

Further, we assessed the diagnostic performance of the LFA device in 50% saliva by evaluating sensitivity and specificity. At 100 nM GP120_MB_, the device achieved a sensitivity of 99%, correctly identifying 100 true positives (TP) with only one false negative (FN). Specificity testing in the absence of GP120_MB_ yielded a 98.02% specificity, with 99 true negatives (TN) and two false positives (FP). To evaluate the potential for off-target signal generation due to cross-reactivity with structurally similar trimeric viral proteins, we tested the device against influenza hemagglutinin (HA) trimers and SARS-CoV-2 spike proteins **(Supplementary Figure S10)**. The test line exhibited strong colorimetric response exclusively in the presence of GP120_MB_, with negligible signal detected for non-HIV protein controls. These results provide compelling evidence of the high specificity and biological compatibility of the DNA-Net_HINT_–based LFA platform for HIV-1 detection.

The DNA-Net_HD4_–based LFA device was also evaluated using the same experimental framework. In this case, DNA-Net_HD4_ constructs were immobilized on the test line (DNA-Net_HD4atT_), and a corresponding AuNS_DNA-Net-HD4_ reporter conjugate was used. Serial dilutions of GP120_MB_ (ranging from 0.1 µM to 1.93 µM) were introduced in running buffer to assess binding affinity. The resulting data revealed that DNA-Net_HD4_ exhibited relatively weak binding to the GP120_MB_ protein, with a *K*_*D*_ of 0.58 ± 0.02 µM **(Supplementary Figures S11A–S11B)**, highlighting the limited affinity of the HD4 aptamer for GP120_MB_. Despite this, the LFA device demonstrated minimal nonspecific interactions when tested across a range of biologically relevant fluids, including saliva, serum, and urine **(Supplementary Figures S11C–S11D)**. Notably, the device maintained operational integrity in a 50% saliva environment, with a *K*_*D*_ of 0.63 ± 0.04 µM **(Supplementary Figures S12A–S12B)**, closely matching the value obtained under aqueous conditions (0% saliva, **Supplementary Figure S11B)**. In terms of target specificity, the DNA-Net_HD4_ LFA device successfully discriminated GP120_MB_ from structurally related trimeric proteins, including influenza hemagglutinin (HA) and the SARS-CoV-2 spike **(Supplementary Figures S12C–S12D)**. However, due to the suboptimal binding performance of the HD4 aptamer in this assay configuration, we limited downstream virus-level testing to the DNA-Net_HINT_ –based device, which had demonstrated superior affinity and diagnostic utility.

### Device performance with HIV-1 viruses and benchmarking against commercial RDTs

One of the objectives of this study was to develop a rapid, reliable, and user-friendly HIV self-testing device capable of detecting low viral loads, particularly during the acute phase of infection. To assess diagnostic performance, the DNA-Net_HINT_–based LFA device was evaluated using two classes of HIV-1 samples: pseudoviruses (PV) and three clinical specimens collected between years 2006–2007. Serial dilutions of each viral sample were prepared in 1× PBS containing 50% saliva to mimic physiologically relevant conditions. For each test, 100 µL of diluted virus sample was applied to the LFA device. A progressive increase in test line intensity was observed with increasing viral load, consistent with the formation of a sandwich complex between the virus, the DNA-Net_HINTatT_ capture probe, and the AuNS_DNA-Net-HINT_ reporter conjugate **(Figures 5a–d)**. The device demonstrated a broad dynamic detection range: from 10^3^ to 10^6^ viral copies for pseudoviruses **(Figure 5a**) and from 10^3^ to 10^5^ viral copies for clinical HIV-1 samples **(Figures 5b-d)**. To benchmark the performance of our device, we conducted a head-to-head comparison using the FDA-approved fourth-generation *Abbott Determine™ HIV-1/2 Ag/Ab Combo* RDT, a widely used commercial product. When tested with clinical samples diluted in 50% saliva, the commercial RDT failed to yield detectable test line signals below 10^5^ viral copies/mL **(Figure 5e)**. In contrast, our device exhibited markedly superior sensitivity. To quantitatively determine the limit of detection (LOD), we applied a four-parameter logistic curve fitting model as previously described^33^. The LODs calculated for the DNA-Net_HINT_–based LFA were 446, 328, 341, and 345 viral copies for the PV and three clinical samples, respectively **(Figure 5f)**. The high analytical sensitivity of our LFA device suggests its potential for early HIV-1 diagnosis during the acute infection phase. This capability is particularly critical for improving linkage to care and minimizing the risk of transmission. Additionally, sensitive and rapid detection plays a pivotal role in clinical decision-making, such as the timely administration of intrapartum prophylaxis and antiretroviral therapy to prevent vertical transmission, thereby contributing meaningfully to public health HIV prevention strategies^34^.

**Figure 5.**
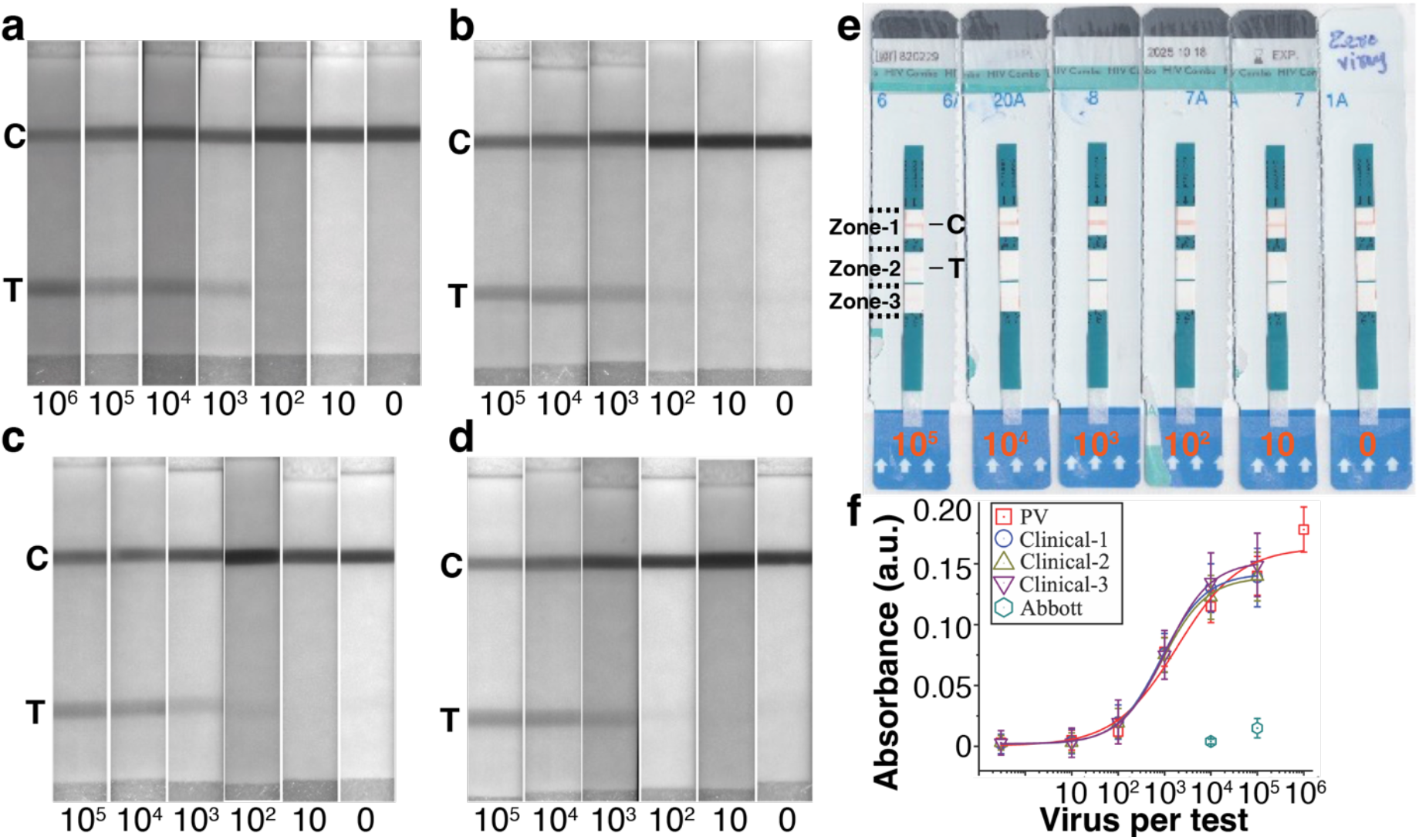
The performance of the LFA device was assessed using HIV virus samples and comparison of device performance with commercially available rapid antigen tests. **(a)** Pseudoviral (PV) and **(b-d)** three clinical samples were systematically diluted (from 10^6^ –10^1^) and (from 10^5^ –10^1^), respectively in 50% saliva environment, while a buffer consisting of 1X PBS and 50% saliva served as the negative control (0 virus). **(e)** Commercial antigen rapid test (Abbott) was utilized with one clinical viral sample. Zone-1: Control zone, Zone-2: Antigen test zone, Zone-3: Antibody test zone, C: Control line, T: Test line. **(f)** Absorbance values from the test line were plotted against the total number of viral particles per test, providing an evaluative measure of sensitivity of our LFA device and commercial rapid antigen tests. The presented data are expressed as the mean ± SD (standard deviation), with n = 3 biologically independent samples.

### Multivalent DNA-Net constructs confer potent antiviral activity against HIV-1

In addition to its diagnostic capabilities, we explored the antiviral potential of the DNA-Net platform by leveraging its multivalent aptamer display to inhibit viral entry. Previous studies have shown that DNA nanostructures can be engineered to engage viral surfaces, thereby blocking infection^21,22,35,36^. In our system, NAMD simulations and SPR analyses revealed that the enhanced binding affinity of DNA-Net constructs stems from the spatial organization of monovalent aptamers into trimeric clusters. This configuration closely mimics the ∼6 nm intra-trimer spacing of GP120 spikes on the HIV-1 envelope, maximizing binding site occupancy and enabling avidity-driven multivalent interactions critical for effective viral neutralization.

To experimentally evaluate this antiviral potential, we conducted flow cytometry–based assays using U87MG cells challenged with DNA-Net-Aptamer treated HIV-1 pseudoviruses and UV-inactivated virions. Viral infection levels were quantified by measuring background-corrected and normalized median fluorescence intensities (MFI). In initial tests, free HD4 and HINT aptamers required micromolar concentrations to achieve 50% inhibition, yielding EC_50_ values of 2.93 ± 0.32 µM and 2.43 ± 0.33 µM, respectively **(Figures S13A–B; Supplementary Tables S3– S4)**. Due to practical limitations in producing DNA-Net constructs at similar micromolar concentrations, we used a 600 nM stock solution and conducted two-fold serial dilutions (1.82– 60 nM). As a negative control, we employed a DNA-Net construct functionalized with a SARS-CoV-2 spike-targeting aptamer^37^ (DNA-Net_RBD_), assembled in the same format as DNA-Net_HINT_.

Remarkably, both DNA-Net_HINT_ and DNA-Net_HD4_ displayed potent viral inhibition at nanomolar (aptamer-equivalent) concentrations in the pseudovirus assay **(Figures S14A, S14C)**. In contrast, only DNA-Net_HINT_ retained modest antiviral activity against UV-inactivated virions **(Figure S14B)**, while DNA-Net_HD4_ failed to show any measurable efficacy **(Figure S14D)**. We also observed reduced infectivity of untreated UV-inactivated virions, which could be attributed to UV-induced conformational disruptions in GP120 structure thus impairing receptor engagement.

Across all tested concentrations in the pseudovirus group, DNA-Net constructs consistently reduced viral loads to below 50% of the uninfected control, while free aptamers showed minimal inhibition, with viral loads comparable to those in untreated or negative control samples **(Figures 6a–c)**. Comparative analysis at the highest concentrations revealed that DNA-Net_HINT_ outperformed both DNA-Net_HD4_ and the respective monomeric aptamers **(Figure 6a)**. Strikingly, DNA-Net_HINT_ achieved an EC_50_ of approximately 1.82 nM, representing greater than ∼1,000-fold enhancement in antiviral potency relative to the free HINT aptamer **(Figure 6c)**.

**Figure 6.**
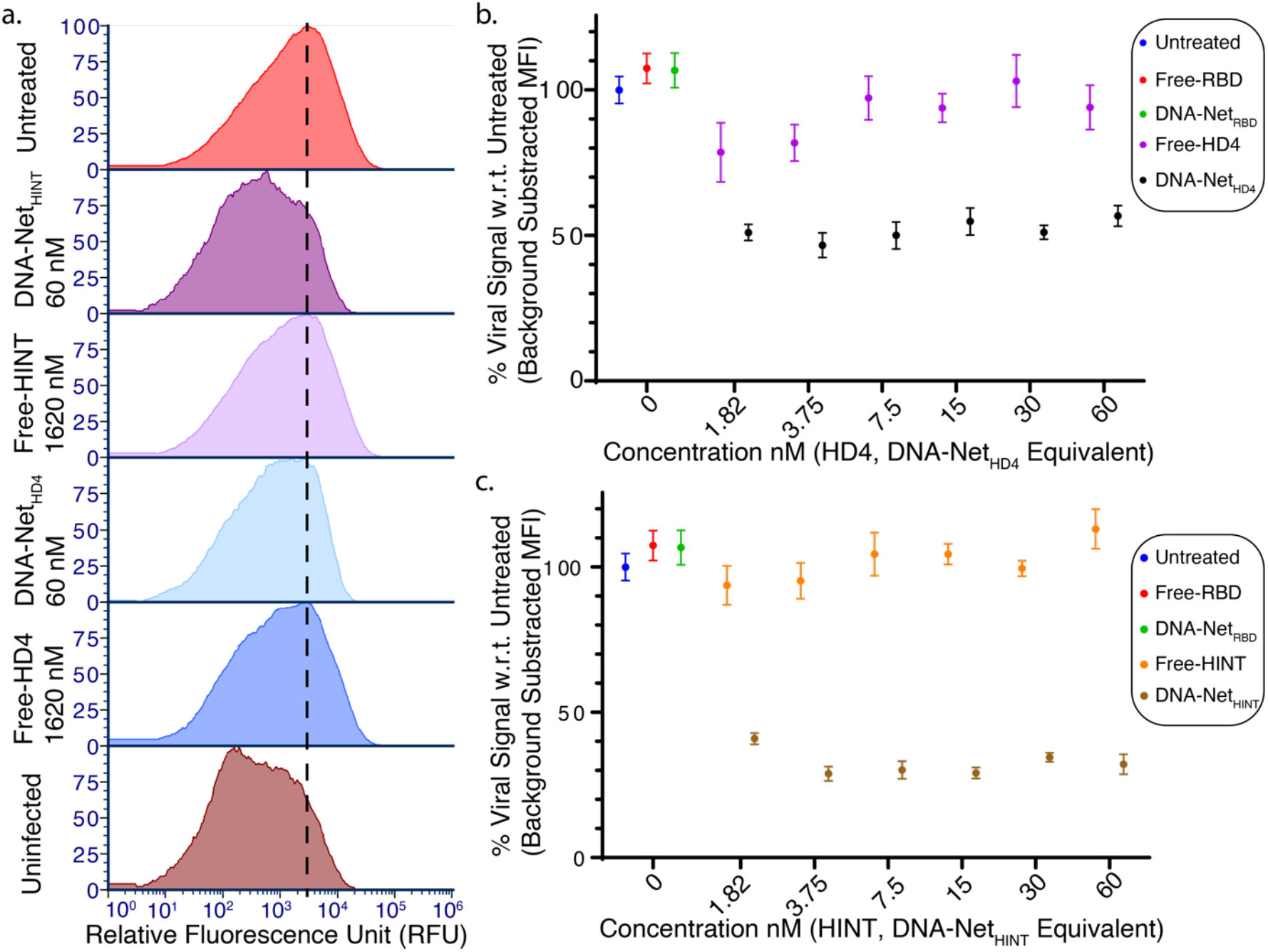
DNA-Net–based aptamer constructs demonstrate enhanced inhibition of active HIV-1 infection in U87MG cells infected with pseudotyped virions (MOI = 1000). **(a)** Flow cytometry analysis comparing concentration-matched free aptamers and DNA-Net–aptamer constructs shows that DNA-Net_HINT_ exhibits significantly improved viral inhibition compared to other constructs and free aptamers at equivalent concentrations. DNA-Net_HINT_ treatment resulted in infection levels approaching those of uninfected controls, suggesting highly effective prevention of viral entry. Serial dilutions of **(b)** DNA-Net_HD4_ and **(c)** DNA-Net_HINT_ were prepared from a 600 nM stock using two-fold serial dilution, yielding a final concentration range of 1.82–60 nM. Free aptamer concentrations were 27-fold higher than their DNA-Net–based counterparts across this range. The untreated virus group (no aptamer or DNA-Net-aptamer constructs) served as a positive infection control, while uninfected cells (no virus added) served as a negative control. DNA-Net_RBD_ (60 nM) and free RBD aptamer^38^ (1620 nM) were included as additional negative controls. Data represent six biologically independent experiments and are presented as mean ± SEM (n = 6).

Taken together, these findings establish that the spatially patterned, multivalent DNA-Net scaffold not only augments target affinity but also confers substantial antiviral efficacy. In addition to functioning as a highly sensitive diagnostic platform, DNA-Net–aptamer constructs exhibit strong potential as modular, programmable antiviral agents for HIV-1 therapeutic applications.

## Conclusions

This study presents a modular DNA nanotechnology–based platform that integrates molecular recognition and antiviral functionality for the broad-spectrum detection and inhibition of HIV-1. Through the rational design of a high-affinity aptamer (HINT) and its spatial presentation on a net-shaped DNA scaffold (DNA-Net), we achieved sub-picomolar binding affinity to trimeric GP120 envelope proteins, validated via molecular dynamics simulations and SPR analyses. When incorporated into a LFA format, the DNA-Net_HINT_ construct enabled ultrasensitive and specific detection of intact HIV-1 virions in biologically relevant conditions such as saliva, achieving a limit of detection as low as 328 viral copies per test, significantly outperforming current fourth-generation commercial RDTs.

Beyond diagnostics, DNA-Net constructs also conferred potent antiviral activity. Flow cytometry– based inhibition assays demonstrated that multivalent spatial patterning of aptamers on the DNA-Net significantly enhanced HIV-1 neutralization, with DNA-Net_HINT_ achieving an EC_50_ of ∼1.82 nM, representing over ∼1,000-fold improvement over the monomeric HINT aptamer with EC_50_ of 2.43 ± 0.33 µM. These findings underscore the important role of nanoscale spatial organization and avidity-driven multivalent binding in improving both diagnostic sensitivity and antiviral efficacy. The modularity of the platform may allow facile adaptation to other enveloped viruses by exchanging aptamer sequences or tuning nanostructure geometry, offering broad utility for pandemic preparedness, point-of-care diagnostics, and targeted antiviral interventions. Overall, this work highlights the transformative potential of designer DNA nanostructures in addressing long-standing challenges in infectious disease detection and prevention.

## Supporting Information

The Supporting Information includes materials, methods, supplementary figures, tables, and references.

## Acknowledgements

The authors acknowledge the support from NIH (R01AI159454 and R21AI166898)

## Data Availability

All the data are included in the submitted manuscript including the Supporting Materials.

## For Table of Contents Only

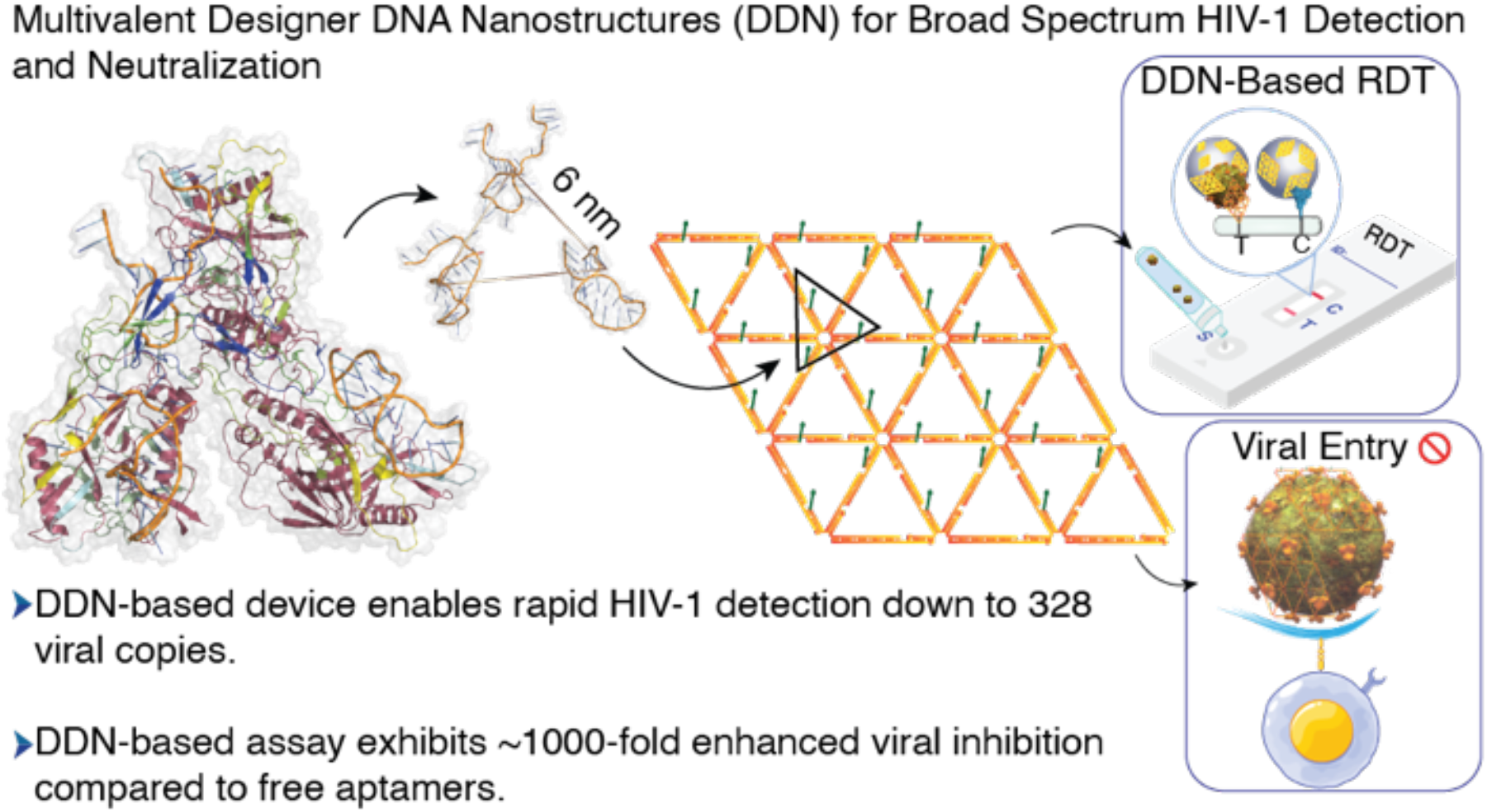

## References

1 van Schalkwyk, C., Mahy, M., Johnson, L. F. & Imai-Eaton, J. W. Updated Data and Methods for the 2023 UNAIDS HIV Estimates. JAIDS Journal of Acquired Immune Deficiency Syndromes 95, e1–e4, doi:10.1097/qai.0000000000003344 (2024).

2 Gulinaizhaer, A., Zou, M., Ma, S., Yao, Y., Fan, X. & Wu, G. Isothermal nucleic acid amplification technology in HIV detection. The Analyst 148, 1189–1208, doi:10.1039/d2an01813f (2023).

3 Fox, J., Dunn, H. & O’Shea, S. Low rates of p24 antigen detection using a fourth-generation point of care HIV test. Sexually Transmitted Infections 87, 178–179, doi:10.1136/sti.2010.042564 (2010).

4 Fenouillet, E., Blanes, N., Coutellier, A. & Gluckman, J. C. Relationship between Anti-p24 Antibody Levels and p24 Antigenemia in HIV-Infected Patients. AIDS Research and Human Retroviruses 9, 1251–1255, doi:10.1089/aid.1993.9.1251 (1993).

5 McMichael, A. J., Borrow, P., Tomaras, G. D., Goonetilleke, N. & Haynes, B. F. The immune response during acute HIV-1 infection: clues for vaccine development. Nature Reviews Immunology 10, 11–23, doi:10.1038/nri2674 (2009).

6 Brenner, Bluma G., Roger, M., Routy, J. P., Moisi, D., Ntemgwa, M., Matte, C., Baril, J. G., Thomas, R., Rouleau, D., Bruneau, J., Leblanc, R., Legault, M., Tremblay, C., Charest, H. & Wainberg, Mark A. High Rates of Forward Transmission Events after Acute/Early HIV-1 Infection. The Journal of infectious diseases 195, 951–959, doi:10.1086/512088 (2007).

7 Pai, N. P., Aghokeng, A. F., Mpoudi-Ngole, E., Dimodi, H., Atem-Tambe, A., Tongo, M., Butel, C., Delaporte, E. & Peeters, M. Inaccurate Diagnosis of HIV-1 Group M and O Is a Key Challenge for Ongoing Universal Access to Antiretroviral Treatment and HIV Prevention in Cameroon. PLoS ONE 4, doi:10.1371/journal.pone.0007702 (2009).

8 Duong, Y. T., Mavengere, Y., Patel, H., Moore, C., Manjengwa, J., Sibandze, D., Rasberry, C., Mlambo, C., Li, Z., Emel, L., Bock, N., Moore, J., Nkambule, R., Justman, J., Reed, J., Bicego, G., Ellenberger, D. L., Nkengasong, J. N., Parekh, B. S. & Caliendo, A. M. Poor Performance of the Determine HIV-1/2 Ag/Ab Combo Fourth-Generation Rapid Test for Detection of Acute Infections in a National Household Survey in Swaziland. Journal of clinical microbiology 52, 3743–3748, doi:10.1128/jcm.01989-14 (2014).

9 Julien, J. P., Cupo, A., Sok, D., Stanfield, R. L., Lyumkis, D., Deller, M. C., Klasse, P. J., Burton, D. R., Sanders, R. W., Moore, J. P., Ward, A. B. & Wilson, I. A. Crystal structure of a soluble cleaved HIV-1 envelope trimer. Science 342, 1477–1483, doi:10.1126/science.1245625 (2013).

10 Zhu, P., Chertova, E., Bess, J., Lifson, J. D., Arthur, L. O., Liu, J., Taylor, K. A. & Roux, K. H. Electron tomography analysis of envelope glycoprotein trimers on HIV and simian immunodeficiency virus virions. Proceedings of the National Academy of Sciences 100, 15812–15817, doi:10.1073/pnas.2634931100 (2003).

11 Ratanabunyong, S., Aeksiri, N., Yanaka, S., Yagi-Utsumi, M., Kato, K., Choowongkomon, K. & Hannongbua, S. Characterization of New DNA Aptamers for Anti-HIV-1 Reverse Transcriptase. Chembiochem 22, 915–923, doi:10.1002/cbic.202000633 (2020).

12 Andreola, M.-L., Pileur, F., Calmels, C., Ventura, M., Tarrago-Litvak, L., Toulmé, J.-J. & Litvak, S. DNA Aptamers Selected against the HIV-1 RNase H Display in Vitro Antiviral Activity. Biochemistry 40, 10087–10094, doi:10.1021/bi0108599 (2001).

13 Liu, L., Tibbs, J., Li, N., Bacon, A., Shepherd, S., Lee, H., Chauhan, N., Demirci, U., Wang, X. & Cunningham, B. T. A photonic resonator interferometric scattering microscope for label-free detection of nanometer-scale objects with digital precision in point-of-use environments. Biosensors and Bioelectronics 228, doi:10.1016/j.bios.2023.115197 (2023).

14 Rizvi, A. S., Murtaza, G., Zhang, W., Xue, M., Qiu, L. & Meng, Z. Aptamer-linked photonic crystal hydrogel sensor for rapid point-of-care detection of human immuno-deficiency virus-1 (HIV-1). Journal of Pharmaceutical and Biomedical Analysis 227, doi:10.1016/j.jpba.2022.115104 (2023).

15 Rizvi, A. S., Murtaza, G., Xu, X., Gao, P., Qiu, L. & Meng, Z. Aptamer-Linked Photonic Crystal Assay for High-Throughput Screening of HIV and SARS-CoV-2. Analytical Chemistry, doi:10.1021/acs.analchem.2c03467 (2022).

16 Njau, B., Ostermann, J., Brown, D., Mühlbacher, A., Reddy, E. & Thielman, N. HIV testing preferences in Tanzania: a qualitative exploration of the importance of confidentiality, accessibility, and quality of service. BMC Public Health 14, doi:10.1186/1471-2458-14-838 (2014).

17 Zhu, P., Liu, J., Bess, J., Jr., Chertova, E., Lifson, J. D., Grise, H., Ofek, G. A., Taylor, K. & Roux, K. H. Distribution and three-dimensional structure of AIDS virus envelope spikes. Nature 441, 847–852, doi:10.1038/nature04817 (2006).

18 Umrao, S., Dwivedy, A. & Wang, X. In Nano-Engineering at Functional Interfaces for Multi-Disciplinary Applications 445–472 (2025).

19 Galimidi, Rachel P., Klein, Joshua S., Politzer, Maria S., Bai, S., Seaman, Michael S., Nussenzweig, Michel C., West, Anthony P. & Bjorkman, Pamela J. Intra-Spike Crosslinking Overcomes Antibody Evasion by HIV-1. Cell 160, 433–446, doi:10.1016/j.cell.2015.01.016 (2015).

20 Rall, G. F., Klein, J. S. & Bjorkman, P. J. Few and Far Between: How HIV May Be Evading Antibody Avidity. PLoS Pathogens 6, doi:10.1371/journal.ppat.1000908 (2010).

21 Chauhan, N., Xiong, Y., Ren, S., Dwivedy, A., Magazine, N., Zhou, L., Jin, X., Zhang, T., Cunningham, B. T., Yao, S., Huang, W. & Wang, X. Net-Shaped DNA Nanostructures Designed for Rapid/Sensitive Detection and Potential Inhibition of the SARS-CoV-2 Virus. Journal of the American Chemical Society 145, 20214–20228, doi:10.1021/jacs.2c04835 (2023).

22 Kwon, P. S., Ren, S., Kwon, S. J., Kizer, M. E., Kuo, L., Xie, M., Zhu, D., Zhou, F., Zhang, F., Kim, D., Fraser, K., Kramer, L. D., Seeman, N. C., Dordick, J. S., Linhardt, R. J., Chao, J. & Wang, X. Designer DNA architecture offers precise and multivalent spatial pattern-recognition for viral sensing and inhibition. Nat Chem 12, 26–35, doi:10.1038/s41557-019-0369-8 (2020).

23 Fischer, A. E., Abrahams, M., Shankland, L., Lalla-Edward, S. T., Edward, V. A. & De Wit, J. The evolution of HIV self-testing and the introduction of digital interventions to improve HIV self-testing. Frontiers in Reproductive Health 5, doi:10.3389/frph.2023.1121478 (2023).

24 Ince, B. & Sezgintürk, M. K. Lateral flow assays for viruses diagnosis: Up-to-date technology and future prospects. TrAC Trends in Analytical Chemistry 157, doi:10.1016/j.trac.2022.116725 (2022).

25 Sepehri Zarandi, H., Behbahani, M. & Mohabatkar, H. In Silico Selection of Gp120 ssDNA Aptamer to HIV-1. SLAS Discovery 25, 1087–1093, doi:10.1177/2472555220923331 (2020).

26 van Zundert, G. C. P., Rodrigues, J. P. G. L. M., Trellet, M., Schmitz, C., Kastritis, P. L., Karaca, E., Melquiond, A. S. J., van Dijk, M., de Vries, S. J. & Bonvin, A. M. J. J. The HADDOCK2.2 Web Server: User-Friendly Integrative Modeling of Biomolecular Complexes. Journal of molecular biology 428, 720–725, doi:10.1016/j.jmb.2015.09.014 (2016).

27 Phillips, J. C., Hardy, D. J., Maia, J. D. C., Stone, J. E., Ribeiro, J. V., Bernardi, R. C., Buch, R., Fiorin, G., Hénin, J., Jiang, W., McGreevy, R., Melo, M. C. R., Radak, B. K., Skeel, R. D., Singharoy, A., Wang, Y., Roux, B., Aksimentiev, A., Luthey-Schulten, Z., Kalé, L. V., Schulten, K., Chipot, C. & Tajkhorshid, E. Scalable molecular dynamics on CPU and GPU architectures with NAMD. The Journal of Chemical Physics 153, doi:10.1063/5.0014475 (2020).

28 Liu, J., Bartesaghi, A., Borgnia, M. J., Sapiro, G. & Subramaniam, S. Molecular architecture of native HIV-1 gp120 trimers. Nature 455, 109–113, doi:10.1038/nature07159 (2008).

29 Wymant, C., Bezemer, D., Blanquart, F., Ferretti, L., Gall, A., Hall, M., Golubchik, T., Bakker, M., Ong, S. H., Zhao, L., Bonsall, D., de Cesare, M., MacIntyre-Cockett, G., Abeler-Dörner, L., Albert, J., Bannert, N., Fellay, J., Grabowski, M. K., Gunsenheimer-Bartmeyer, B., Günthard, H. F., Kivelä, P., Kouyos, R. D., Laeyendecker, O., Meyer, L., Porter, K., Ristola, M., van Sighem, A., Berkhout, B., Kellam, P., Cornelissen, M., Reiss, P., Fraser, C., Aubert, V., Battegay, M., Bernasconi, E., Böni, J., Braun, D. L., Bucher, H. C., Burton-Jeangros, C., Calmy, A., Cavassini, M., Dollenmaier, G., Egger, M., Elzi, L., Fehr, J., Fellay, J., Furrer, H., Fux, C. A., Gorgievski, M., Günthard, H., Haerry, D., Hasse, B., Hirsch, H. H., Hoffmann, M., Hösli, I., Kahlert, C., Kaiser, L., Keiser, O., Klimkait, T., Kouyos, R., Kovari, H., Ledergerber, B., Martinetti, G., de Tejada, B. M., Marzolini, C., Metzner, K., Müller, N., Nadal, D., Nicca, D., Pantaleo, G., Rauch, A., Regenass, S., Rudin, C., Schöni-Affolter, F., Schmid, P., Speck, R., Stöckle, M., Tarr, P., Trkola, A., Vernazza, P., Weber, R., Yerly, S., van der Valk, M., Geerlings, S. E., Goorhuis, A., Hovius, J. W., Lempkes, B., Nellen, F. J. B., van der Poll, T., Prins, J. M., Reiss, P., van Vugt, M., Wiersinga, W. J., Wit, F. W. M. N., van Duinen, M., van Eden, J., Hazenberg, A., van Hes, A. M. H., Pijnappel, F. J. J., Smalhout, S. Y., Weijsenfeld, A. M., Jurriaans, S., Back, N. K. T., Zaaijer, H. L., Berkhout, B., Cornelissen, M. T. E., Schinkel, C. J., Wolthers, K. C., Peters, E. J. G., van Agtmael, M. A., Autar, R. S., Bomers, M., Sigaloff, K. C. E., Heitmuller, M., Laan, L. M., Ang, C. W., van Houdt, R., Jonges, M., Kuijpers, T. W., Pajkrt, D., Scherpbier, H. J., de Boer, C., van der Plas, A., van den Berge, M., Stegeman, A., Baas, S., Hage de Looff, L., Buiting, A., Reuwer, A., Veenemans, J., Wintermans, B., Pronk, M. J. H., Ammerlaan, H. S. M., van den Bersselaar, D. N. J., de Munnik, E. S., Deiman, B., Jansz, A. R., Scharnhorst, V., Tjhie, J., Wegdam, M. C. A., van Eeden, A., Nellen, J., Brokking, W., Elsenburg, L. J. M., Nobel, H., van Kasteren, M. E. E., Berrevoets, M. A. H., Brouwer, A. E., Adams, A., van Erve, R., de Kruijf-van de Wiel, B. A. F. M., Keelan-Phaf, S., van de Ven, B., van der Ven, B., Buiting, A. G. M., Murck, J. L., de Vries-Sluijs, T. E. M. S., Bax, H. I., van Gorp, E. C. M., de Jong-Peltenburg, N. C., de MendonÇa Melo, M., van Nood, E., Nouwen, J. L., Rijnders, B. J. A., Rokx, C., Schurink, C. A. M., Slobbe, L., Verbon, A., Bassant, N., van Beek, J. E. A., Vriesde, M., van Zonneveld, L. M., de Groot, J., Boucher, C. A. B., Koopmans, M. P. G., van Kampen, J. J. A., Fraaij, P. L. A., van Rossum, A. M. C., Vermont, C. L., van der Knaap, L. C., Visser, E., Branger, J., Douma, R. A., Cents-Bosma, A. S., Duijf-van de Ven, C. J. H. M., Schippers, E. F., van Nieuwkoop, C., van Ijperen, J. M., Geilings, J., van der Hut, G., van Burgel, N. D., Leyten, E. M. S., Gelinck, L. B. S., Mollema, F., Davids-Veldhuis, S., Tearno, C., Wildenbeest, G. S., Heikens, E., Groeneveld, P. H. P., Bouwhuis, J. W., Lammers, A. J. J., Kraan, S., van Hulzen, A. G. W., Kruiper, M. S. M., van der Bliek, G. L., Bor, P. C. J., Debast, S. B., Wagenvoort, G. H. J., Kroon, F. P., de Boer, M. G. J., Jolink, H., Lambregts, M. M. C., Roukens, A. H. E., Scheper, H., Dorama, W., van Holten, N., Claas, E. C. J., Wessels, E., den Hollander, J. G., El Moussaoui, R., Pogany, K., Brouwer, C. J., Smit, J. V., Struik-Kalkman, D., van Niekerk, T., Pontesilli, O., Lowe, S. H., Oude Lashof, A. M. L., Posthouwer, D., van Wolfswinkel, M. E., Ackens, R. P., Burgers, K., Schippers, J., Weijenberg-Maes, B., van Loo, I. H. M., Havenith, T. R. A., van Vonderen, M. G. A., Kampschreur, L. M., Faber, S., Steeman-Bouma, R., Al Moujahid, A., Kootstra, G. J., Delsing, C. E., van der Burg-van de Plas, M., Scheiberlich, L., Kortmann, W., van Twillert, G., Renckens, R., Ruiter-Pronk, D., van Truijen-Oud, F. A., Cohen Stuart, J. W. T., Jansen, E. R., Hoogewerf, M., Rozemeijer, W., van der Reijden, W. A., Sinnige, J. C., Brinkman, K., van den Berk, G. E. L., Blok, W. L., Lettinga, K. D., de Regt, M., Schouten, W. E. M., Stalenhoef, J. E., Veenstra, J., Vrouenraets, S. M. E., Blaauw, H., Geerders, G. F., Kleene, M. J., Kok, M., Knapen, M., van der Meché, I. B., Mulder-Seeleman, E., Toonen, A. J. M., Wijnands, S., Wttewaal, E., Kwa, D., van Crevel, R., van Aerde, K., Dofferhoff, A. S. M., Henriet, S. S. V., ter Hofstede, H. J. M., Hoogerwerf, J., Keuter, M., Richel, O., Albers, M., Grintjes-Huisman, K. J. T., de Haan, M., Marneef, M., Strik-Albers, R., Rahamat-Langendoen, J., Stelma, F. F., Burger, D., Gisolf, E. H., Hassing, R. J., Claassen, M., ter Beest, G., van Bentum, P. H. M., Langebeek, N., Tiemessen, R., Swanink, C. M. A., van Lelyveld, S. F. L., Soetekouw, R., van der Prijt, L. M. M., van der Swaluw, J., Bermon, N., van der Reijden, W. A., Jansen, R., Herpers, B. L., Veenendaal, D., Verhagen, D. W. M., Lauw, F. N., van Broekhuizen, M. C., van Wijk, M., Bierman, W. F. W., Bakker, M., Kleinnijenhuis, J., Kloeze, E., Middel, A., Postma, D. F., Schölvinck, E. H., Stienstra, Y., Verhage, A. R., Wouthuyzen-Bakker, M., Boonstra, A., de Groot-de Jonge, H., van der Meulen, P. A., de Weerd, D. A., Niesters, H. G. M., van Leer-Buter, C. C., Knoester, M., Hoepelman, A. I. M., Arends, J. E., Barth, R. E., Bruns, A. H. W., Ellerbroek, P. M., Mudrikova, T., Oosterheert, J. J., Schadd, E. M., van Welzen, B. J., Aarsman, K., Griffioen-van Santen, B. M. G., de Kroon, I., van Berkel, M., van Rooijen, C. S. A. M., Schuurman, R., Verduyn-Lunel, F., Wensing, A. M. J., Bont, L. J., Geelen, S. P. M., Loeffen, Y. G. T., Wolfs, T. F. W., Nauta, N., Rooijakkers, E. O. W., Holtsema, H., Voigt, R., van de Wetering, D., Alberto, A., van der Meer, I., Rosingh, A., Halaby, T., Zaheri, S., Boyd, A. C., Bezemer, D. O., van Sighem, A. I., Smit, C., Hillebregt, M., de Jong, A., Woudstra, T., Bergsma, D., Meijering, R., van de Sande, L., Rutkens, T., van der Vliet, S., de Groot, L., van den Akker, M., Bakker, Y., El Berkaoui, A., Bezemer, M., Brétin, N., Djoechro, E., Groters, M., Kruijne, E., Lelivelt, K. J., Lodewijk, C., Lucas, E., Munjishvili, L., Paling, F., Peeck, B., Ree, C., Regtop, R., Ruijs, Y., Schoorl, M., Schnörr, P., Scheigrond, A., Tuijn, E., Veenenberg, L., Visser, K. M., Witte, E. C., Ruijs, Y., Van Frankenhuijsen, M., Allegre, T., Makhloufi, D., Livrozet, J.-M., Chiarello, P., Godinot, M., Brunel-Dalmas, F., Gibert, S., Trepo, C., Peyramond, D., Miailhes, P., Koffi, J., Thoirain, V., Brochier, C., Baudry, T., Pailhes, S., Lafeuillade, A., Philip, G., Hittinger, G., Assi, A., Lambry, V., Rosenthal, E., Naqvi, A., Dunais, B., Cua, E., Pradier, C., Durant, J., Joulie, A., Quinsat, D., Tempesta, S., Ravaux, I., Martin, I. P., Faucher, O., Cloarec, N., Champagne, H., Pichancourt, G., Morlat, P., Pistone, T., Bonnet, F., Mercie, P., Faure, I., Hessamfar, M., Malvy, D., Lacoste, D., Pertusa, M.-C., Vandenhende, M.-A., Bernard, N., Paccalin, F., Martell, C., Roger-Schmelz, J., Receveur, M.-C., Duffau, P., Dondia, D., Ribeiro, E., Caltado, S., Neau, D., Dupont, M., Dutronc, H., Dauchy, F., Cazanave, C., Vareil, M.-O., Wirth, G., Le Puil, S., Pellegrin, J.-L., Raymond, I., Viallard, J.-F., Chaigne de Lalande, S., Garipuy, D., Delobel, P., Obadia, M., Cuzin, L., Alvarez, M., Biezunski, N., Porte, L., Massip, P., Debard, A., Balsarin, F., Lagarrigue, M., Prevoteau du Clary, F., Aquilina, C., Reynes, J., Baillat, V., Merle, C., Lemoing, V., Atoui, N., Makinson, A., Jacquet, J. M., Psomas, C., Tramoni, C., Aumaitre, H., Saada, M., Medus, M., Malet, M., Eden, A., Neuville, S., Ferreyra, M., Sotto, A., Barbuat, C., Rouanet, I., Leureillard, D., Mauboussin, J.-M., Lechiche, C., Donsesco, R., Cabie, A., Abel, S., Pierre-Francois, S., Batala, A.-S., Cerland, C., Rangom, C., Theresine, N., Hoen, B., Lamaury, I., Fabre, I., Schepers, K., Curlier, E., Ouissa, R., Gaud, C., Ricaud, C., Rodet, R., Wartel, G., Sautron, C., Beck-Wirth, G., Michel, C., Beck, C., Halna, J.-M., Kowalczyk, J., Benomar, M., Drobacheff-Thiebaut, C., Chirouze, C., Faucher, J.-F., Parcelier, F., Foltzer, A., Haffner-Mauvais, C., Hustache Mathieu, M., Proust, A., Piroth, L., Chavanet, P., Duong, M., Buisson, M., Waldner, A., Mahy, S., Gohier, S., Croisier, D., May, T., Delestan, M., Andre, M., Zadeh, M. M., Martinot, M., Rosolen, B., Pachart, A., Martha, B., Jeunet, N., Rey, D., Cheneau, C., Partisani, M., Priester, M., Bernard-Henry, C., Batard, M.-L., Fischer, P., Berger, J.-L., Kmiec, I., Robineau, O., Huleux, T., Ajana, F., Alcaraz, I., Allienne, C., Baclet, V., Meybeck, A., Valette, M., Viget, N., Aissi, E., Biekre, R., Cornavin, P., Merrien, D., Seghezzi, J.-C., Machado, M., Diab, G., Raffi, F., Bonnet, B., Allavena, C., Grossi, O., Reliquet, V., Billaud, E., Brunet, C., Bouchez, S., Morineau-Le Houssine, P., Sauser, F., Boutoille, D., Besnier, M., Hue, H., Hall, N., Brosseau, D., Souala, F., Michelet, C., Tattevin, P., Arvieux, C., Revest, M., Leroy, H., Chapplain, J.-M., Dupont, M., Fily, F., Patra-Delo, S., Lefeuvre, C., Bernard, L., Bastides, F., Nau, P., Verdon, R., de la Blanchardiere, A., Martin, A., Feret, P., Geffray, L., Daniel, C., Rohan, J., Fialaire, P., Chennebault, J. M., Rabier, V., Abgueguen, P., Rehaiem, S., Luycx, O., Niault, M., Moreau, P., Poinsignon, Y., Goussef, M., Mouton-Rioux, V., Houlbert, D., Alvarez-Huve, S., Barbe, F., Haret, S., Perre, P., Leantez-Nainville, S., Esnault, J.-L., Guimard, T., Suaud, I., Girard, J.-J., Simonet, V., Debab, Y., Schmit, J.-L., Jacomet, C., Weinberck, P., Genet, C., Pinet, P., Ducroix, S., Durox, H., Denes, É., Abraham, B., Gourdon, F., Antoniotti, O., Molina, J.-M., Ferret, S., Lascoux-Combe, C., Lafaurie, M., Colin de Verdiere, N., Ponscarme, D., De Castro, N., Aslan, A., Rozenbaum, W., Pintado, C., Clavel, F., Taulera, O., Gatey, C., Munier, A.-L., Gazaigne, S., Penot, P., Conort, G., Lerolle, N., Leplatois, A., Balausine, S., Delgado, J., Timsit, J., Tabet, M., Gerard, L., Girard, P.-M., Picard, O., Tredup, J., Bollens, D., Valin, N., Campa, P., Bottero, J., Lefebvre, B., Tourneur, M., Fonquernie, L., Wemmert, C., Lagneau, J.-L., Yazdanpanah, Y., Phung, B., Pinto, A., Vallois, D., Cabras, O., Louni, F., Pialoux, G., Lyavanc, T., Berrebi, V., Chas, J., Lenagat, S., Rami, A., Diemer, M., Parrinello, M., Depond, A., Salmon, D., Guillevin, L., Tahi, T., Belarbi, L., Loulergue, P., Zak Dit Zbar, O., Launay, O., Silbermann, B., Leport, C., Alagna, L., Pietri, M.-P., Simon, A., Bonmarchand, M., Amirat, N., Pichon, F., Kirstetter, M., Katlama, C., Valantin, M. A., Tubiana, R., Caby, F., Schneider, L., Ktorza, N., Calin, R., Merlet, A., Ben Abdallah, S., Weiss, L., Buisson, M., Batisse, D., Karmochine, M., Pavie, J., Minozzi, C., Jayle, D., Castel, P., Derouineau, J., Kousignan, P., Eliazevitch, M., Pierre, I., Collias, L., Viard, J.-P., Gilquin, J., Sobel, A., Slama, L., Ghosn, J., Hadacek, B., Thu-Huyn, N., Nait-Ighil, L., Cros, A., Maignan, A., Duvivier, C., Consigny, P. H., Lanternier, F., Shoai-Tehrani, M., Touam, F., Jerbi, S., Bodard, L., Jung, C., Goujard, C., Quertainmont, Y., Duracinsky, M., Segeral, O., Blanc, A., Peretti, D., Cheret, A., Chantalat, C., Dulucq, M. J., Levy, Y., Lelievre, J. D., Lascaux, A. S., Dumont, C., Boue, F., Chambrin, V., Abgrall, S., Kansau, I., Raho-Moussa, M., De Truchis, P., Dinh, A., Davido, B., Marigot, D., Berthe, H., Devidas, A., Chevojon, P., Chabrol, A., Agher, N., Lemercier, Y., Chaix, F., Turpault, I., Bouchaud, O., Honore, P., Rouveix, E., Reimann, E., Belan, A. G., Godin Collet, C., Souak, S., Mortier, E., Bloch, M., Simonpoli, A.-M., Manceron, V., Cahitte, I., Hiraux, E., Lafon, E., Cordonnier, F., Zeng, A.-f., Zucman, D., Majerholc, C., Bornarel, D., Uludag, A., Gellen-Dautremer, J., Lefort, A., Bazin, C., Daneluzzi, V., Gerbe, J., Jeantils, V., Coupard, M., Patey, O., Bantsimba, J., Delllion, S., Paz, P. C., Cazenave, B., Richier, L., Garrait, V., Delacroix, I., Elharrar, B., Vittecoq, D., Bolliot, C., Lepretre, A., Genet, P., Masse, V., Perrone, V., Boussard, J.-L., Chardon, P., Froguel, E., Simon, P., Tassi, S., Avettand Fenoel, V., Barin, F., Bourgeois, C., Cardon, F., Chaix, M.-L., Delfraissy, J. F., Essat, A., Fischer, H., Lecuroux, C., Meyer, L., Petrov-Sanchez, V., Rouzioux, C., Saez-Cirion, A., Seng, R., Kuldanek, K., Mullaney, S., Young, C., Zucchetti, A., Bevan, M.-A., McKernan, S., Wandolo, E., Richardson, C., Youssef, E., Green, P., Faulkner, S., Faville, R., Herman, S., Care, C., Blackman, H., Bellenger, K., Fairbrother, K., Phillips, A., Babiker, A., Delpech, V., Fidler, S., Clarke, M., Fox, J., Gilson, R., Goldberg, D., Hawkins, D., Johnson, A., Johnson, M., McLean, K., Nastouli, E., Post, F., Kennedy, N., Pritchard, J., Andrady, U., Rajda, N., Donnelly, C., McKernan, S., Drake, S., Gilleran, G., White, D., Ross, J., Harding, J., Faville, R., Sweeney, J., Flegg, P., Toomer, S., Wilding, H., Woodward, R., Dean, G., Richardson, C., Perry, N., Gompels, M., Jennings, L., Bansaal, D., Browing, M., Connolly, L., Stanley, B., Estreich, S., Magdy, A., O’Mahony, C., Fraser, P., Jebakumar, S. P. R., David, L., Mette, R., Summerfield, H., Evans, M., White, C., Robertson, R., Lean, C., Morris, S., Winter, A., Faulkner, S., Goorney, B., Howard, L., Fairley, I., Stemp, C., Short, L., Gomez, M., Young, F., Roberts, M., Green, S., Sivakumar, K., Minton, J., Siminoni, A., Calderwood, J., Greenhough, D., DeSouza, C., Muthern, L., Orkin, C., Murphy, S., Truvedi, M., McLean, K., Hawkins, D., Higgs, C., Moyes, A., Antonucci, S., McCormack, S., Lynn, W., Bevan, M., Fox, J., Teague, A., Anderson, J., Mguni, S., Post, F., Campbell, L., Mazhude, C., Russell, H., Gilson, R., Carrick, G., Ainsworth, J., Waters, A., Byrne, P., Johnson, M., Fidler, S., Kuldanek, K., Mullaney, S., Lawlor, V., Melville, R., Sukthankar, A., Thorpe, S., Murphy, C., Wilkins, E., Ahmad, S., Green, P., Tayal, S., Ong, E., Meaden, J., Riddell, L., Loay, D., Peacock, K., Blackman, H., Harindra, V., Saeed, A. M., Allen, S., Natarajan, U., Williams, O., Lacey, H., Care, C., Bowman, C., Herman, S., Devendra, S. V., Wither, J., Bridgwood, A., Singh, G., Bushby, S., Kellock, D., Young, S., Rooney, G., Snart, B., Currie, J., Fitzgerald, M., Arumainayyagam, J., Chandramani, S., Rajamanoharan, S., Robinson, T., Taylor, B., Brewer, C., Mayr, C., Schmidt, W., Speidel, A., Strohbach, F., Arastéh, K., Cordes, C., Stündel, M., Claus, J., Baumgarten, A., Carganico, A., Ingiliz, P., Dupke, S., Freiwald, M., Rausch, M., Moll, A., Schleehauf, D., Hintsche, B., Klausen, G., Jessen, H., Jessen, A., Köppe, S., Kreckel, P., Schranz, D., Fischer, K., Schulbin, H., Speer, M., Glaunsinger, T., Wicke, T., Bieniek, B., Hillenbrand, H., Schlote, F., Lauenroth-Mai, E., Schuler, C., Schürmann, D., Wesselmann, H., Brockmeyer, N., Gehring, P., Schmalöer, D., Hower, M., Spornraft-Ragaller, P., Häussinger, D., Reuter, S., Esser, S., Markus, R., Kreft, B., Berzow, D., Christl, A., Meyer, A., Plettenberg, A., Stoehr, A., Graefe, K., Lorenzen, T., Adam, A., Schewe, K., Weitner, L., Fenske, S., Hansen, S., Stellbrink, H.-J., Wiemer, D., Hertling, S., Schmidt, R., Arbter, P., Claus, B., Galle, P., Jäger, H., Jägel-Guedes, E., Postel, N., Fröschl, M., Spinner, C., Bogner, J., Salzberger, B., Schölmerich, J., Audebert, F., Marquardt, T., Schaffert, A., Schnaitmann, E., Trein, A., Frietsch, B., Müller, M., Ulmer, A., Detering-Hübner, B., Kern, P., Schubert, F., Dehn, G., Schreiber, M., Güler, C., Gunsenheimer-Bartmeyer, B., Schmidt, D., Meixenberger, K. & Bannert, N. A highly virulent variant of HIV-1 circulating in the Netherlands. Science 375, 540–545, doi:10.1126/science.abk1688 (2022).

30 Umrao, S., Zheng, M., Jin, X., Yao, S. & Wang, X. Net-Shaped DNA Nanostructure-Based Lateral Flow Assays for Rapid and Sensitive SARS-CoV-2 Detection. Analytical Chemistry 96, 3291–3299, doi:10.1021/acs.analchem.3c03698 (2024).

31 Gonzalez-Moa, M. J., Van Dorst, B., Lagatie, O., Verheyen, A., Stuyver, L. & Biamonte, M. Proof-of-Concept Rapid Diagnostic Test for Onchocerciasis: Exploring Peptide Biomarkers and the Use of Gold Nanoshells as Reporter Nanoparticles. ACS infectious diseases 4, 912–917, doi:10.1021/acsinfecdis.8b00031 (2018).

32 Ibitoye, M., Frasca, T., Giguere, R. & Carballo-Diéguez, A. Home Testing Past, Present and Future: Lessons Learned and Implications for HIV Home Tests. AIDS and Behavior 18, 933–949, doi:10.1007/s10461-013-0668-9 (2013).

33 Holstein, C. A., Griffin, M., Hong, J. & Sampson, P. D. Statistical Method for Determining and Comparing Limits of Detection of Bioassays. Analytical Chemistry 87, 9795–9801, doi:10.1021/acs.analchem.5b02082 (2015).

34 HIV Testing and Prophylaxis to Prevent Mother-to-Child Transmission in the United States. Pediatrics 122, 1127–1134, doi:10.1542/peds.2008-2175 (2008).

35 Ren, S., Fraser, K., Kuo, L., Chauhan, N., Adrian, A. T., Zhang, F., Linhardt, R. J., Kwon, P. S. & Wang, X. Designer DNA nanostructures for viral inhibition. Nature Protocols 17, 282–326, doi:10.1038/s41596-021-00641-y (2022).

36 Zhou, L., Xiong, Y., Dwivedy, A., Zheng, M., Cooper, L., Shepherd, S., Song, T., Hong, W., Le, L. T. P., Chen, X., Umrao, S., Rong, L., Wang, T., Cunningham, B. T. & Wang, X. Bioinspired designer DNA NanoGripper for virus sensing and potential inhibition. Science Robotics 9, eadi2084, doi:10.1126/scirobotics.adi2084 (2024).

37 Song, Y., Song, J., Wei, X., Huang, M., Sun, M., Zhu, L., Lin, B., Shen, H., Zhu, Z. & Yang, C. Discovery of Aptamers Targeting the Receptor-Binding Domain of the SARS-CoV-2 Spike Glycoprotein. Anal Chem 92, 9895–9900, doi:10.1021/acs.analchem.0c01394 (2020).

